# Assembling an illustrated family-level tree of life for exploration in mobile devices

**DOI:** 10.1101/2021.08.04.454988

**Authors:** Andrés A. Del Risco, Diego A. Chacón, Lucia Ángel, David A. García

## Abstract

Since the concept of the tree of life was introduced by Darwin about a century and a half ago, a considerable fraction of the scientific community has focused its efforts on its reconstruction, with remarkable progress during the last two decades with the advent of DNA sequences. However, the assemblage of a comprehensive tree of life for its exploration has been a difficult task to achieve due to two main obstacles: i) information is scattered into a plethora of individual sources and ii) practical visualization tools for exceptionally large trees are lacking. To overcome both challenges, we aimed to synthetize a family-level tree of life by compiling over 1400 published phylogenetic studies, ensuring that the source trees represent the best phylogenetic hypotheses to date based on a set of objective criteria. Moreover, we dated the synthetic tree by employing over 550 secondary-calibration points, using publicly available sequences for more than 5000 taxa, and by incorporating age ranges from the fossil record for over 2800 taxa. Additionally, we developed a mobile app (Tree of Life) for smartphones in order to facilitate the visualization and interactive exploration of the resulting tree. Interactive features include an easy exploration by zooming and panning gestures of touch screens, collapsing branches, visualizing specific clades as subtrees, a search engine, a timescale to determine extinction and divergence dates, and quick links to Wikipedia. Small illustrations of organisms are displayed at the tips of the branches, to better visualize the morphological diversity of life on earth. Our assembled Tree of Life currently includes over 7000 taxonomic families (about half of the total family-level diversity) and its content will be gradually expanded through regular updates to cover all life on earth at family-level.

## INTRODUCTION

The fact that all life on Earth is connected through common ancestry across a single genealogical tree has fascinated scientists for over one and a half century (Darwin, 1859). Since then, reconstructing the tree of life has become one of the most challenging goals in the biological sciences, prompting the emergence of phylogenetics as a fundamental subdiscipline of evolutionary biology.

Even though the scientific community has made substantial advances towards this goal, especially during the last two decades following the advent of molecular systematics (Pennisi 2013), two major obstacles still stand tall in the way of providing a unified and comprehensive tree of life to browse through the evolutionary relatedness of all life on Earth. Firstly, phylogenetic reconstructions are usually focused on very small fractions of the whole tree of life, so the available fragments of information are widely scattered across thousands of scientific publications which in turn are buried in the often paywalled primary literature. Secondly, due to the high number of terminal branches, having a fully assembled tree of life would require effective and practical ways to navigate and visualize a phylogenetic diagram of such large dimensions.

Some important efforts have been undertaken to tackle both obstacles (although separately) with varying degrees of success. Early projects aiming to reconstruct the tree of life involved the compilation of phylogenies of specific groups of taxa into sectional, hyperlinked webpages (Maddison et al. 2007) or books (Lecointre & Le Guyader 2006; Hedges & Kumar 2009), but these were eventually superseded by the most comprehensive initiative to synthesize phylogenetic and taxonomic information into a single source tree: The Open Tree of Life (hereafter, OTOL: Hinchliff et al. 2015). Although the establishment of the OTOL project represents a huge step in overcoming the first obstacle by the extensive compilation of phylogenies, it is not devoid of shortcomings. Firstly, the construction of its constantly updating synthetic tree mainly depends upon published phylogenies in the form of tree files, which are acquired from either public data repositories or by direct submission by authors. Unfortunately, less than a fifth of the studies provide such files (Drew 2013), meaning that most of the potentially valuable phylogenetic information that is presented in the form of figures in published articles are yet to be incorporated into its synthetic tree. Secondly, the OTOL synthetic tree does not include the temporal component (time-proportional branch lengths) of the Tree of Life, so this information must be consulted elsewhere (e.j., Kumar et al. 2017). Finally, the OTOL project currently aims to reconstruct the evolutionary relationships among living taxa, so extinct taxa known from the vast fossil record has yet to be incorporated into the synthesis.

On the other hand, while many programs that allow users to visualize and navigate tree files have been available for over a decade (Page 2011), only two recently developed web-based tools have attained the goal of providing a complete visualization and exploration of large phylogenies: Onezoom (Rosindell & Harmon 2012) and LifeMap (de Vienne 2016). Both allow users to smoothly explore the tree of life by employing zooming features, and by being also available in the form of mobile apps, they take advantage of touch screens for a more intuitive exploration. These tools were solely developed for visualization, and hence depend on external datasets to explore a given phylogeny, relying on the synthetic tree compiled by the OTOL or the NCBI taxonomic tree. The main limitation of these approaches is the visual formats in which trees are presented; even though they are aesthetically appealing, they greatly deviate from the conventional diagrams in which phylogenies are presented (cladograms, phylograms, timetrees, etc.) and thus hinder the proper inference of the topological relationships depicted in the tree. Additionally, another consequence of these distinctive visualization formats is their inability to incorporate a temporal dimension in the form of branch lengths, which is necessary to determine extinction and divergence dates.

With the aim of overcoming both obstacles while avoiding some of the mentioned pitfalls of previous projects, we introduce the Tree of Life App: a novel tool to visualize and interactively explore a thoroughly compiled, time-calibrated family level-tree of life through a mobile app. We aim to establish this tool as a reference resource for scientific consultation, educational purposes, and even casual exploration for the general public. Tree of Life App is publicly available in the Google Play Store and iPhone AppStore, and it is constantly being updated as new phylogenetic studies are published.

## METHODS

The general procedure for the assemblage of the family-level tree of life consisted mainly in a tree grafting approach, in which higher-level phylogenies are pasted into lower-level phylogenies. The whole procedure, from the initial lists of taxa to the time-calibrated tree file, is depicted in Figure 1 and is thoroughly explained in the sections below.

**Figure 1.**
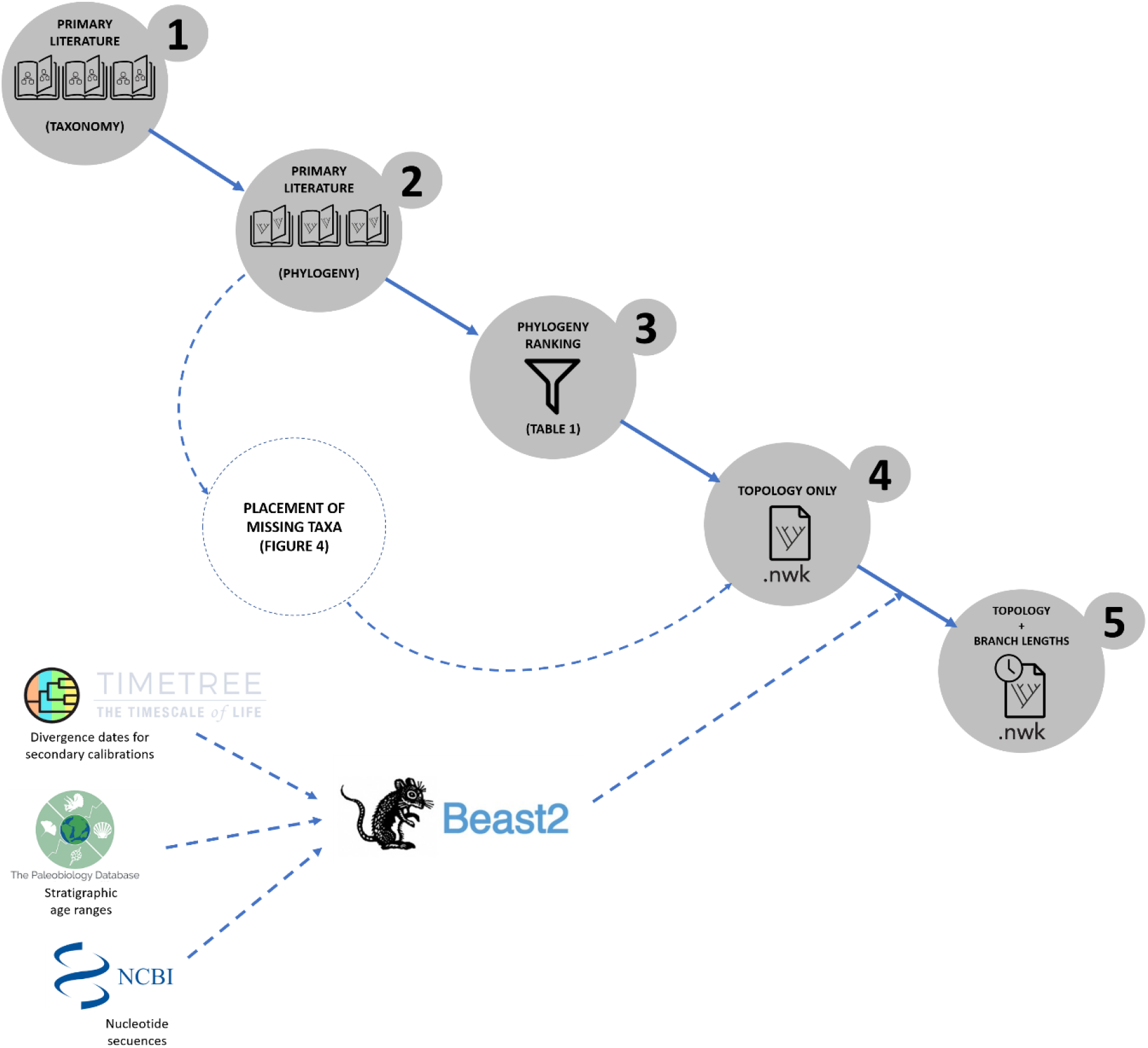
Summary workflow of the tree assembling process. Major steps are enclosed in gray circles, which encompass (1) the establishment of the taxonomic backbones from the primary literature, (2) the obtention of phylogenetic studies, (3) ranking phylogenetic hypotheses given a set of criteria to select the most suitable trees, (4) the transcription of the selected higher ranked phylogenies (and the placement of missing taxa from others) to a Newick file, and (5) the assignation of branch lengths based on a comprehensive tree-dating analysis with BEAST2, resulting in the final synthesized tree.

### Taxonomic Input

To assemble the family-level tree of life, the first step was establishing comprehensive taxonomic lists for large groups of organisms based on clade-specific, published taxonomic backbones covering at least family-ranked taxa. We employed the comprehensive family-level classification of living organisms of Ruggiero (2014) as a starting point and as the default taxonomic backbone for groups without specialized proposals. Although different taxonomic proposals or treatments may exist for a given group, we chose what we consider to be the best taxonomic backbone based on their congruence with current phylogenetic hypotheses (i.e., recognition of mostly monophyletic families). Each family-level taxonomic backbone is then cited in and displayed in the secondary information window of its corresponding supra-familial taxon. For any given group, a taxonomic family list may or may not be a composite of more than one source. Such is the case for groups that have a rich fossil record, as are most major vertebrate and arthropod clades, among others. In these cases, taxonomic lists are usually composed of a source for extant families, and another one for extinct families (which are often excluded in the former). For most groups, fossil families are taken from the PaleoBiology Database (hereafter PBDB; http://paleobiodb.org) taxonomy, unless otherwise specified in secondary information windows. Taxa that are currently unassigned to a family-level group, such as genera *incertae sedis* or fossil taxa that do not always employ Linnaean ranks are not included in the procedure.

### Phylogenetic arrangements

The second step in the procedure is arranging taxa into cladograms by manually writing text files in Newick tree notation. These topological arrangements are direct transcriptions of phylogenies published as figures in the primary literature that were identified and collected after a comprehensive search with the Google Scholar engine. By using figures rather than tree files, we avoid the overdependence of available datasets in public repositories, which are lacking for over 80% of the published phylogenetic knowledge (Drew et al. 2013). Family-level phylogenies were separately built in this way for lower-level clades and were then progressively condensed into larger and higher-level trees, whose backbone topologies were inferred and manually written following the same approach described previously. Because the operational taxonomic units (OTUs) of published phylogenies are mostly species-level, terminals are tagged with its corresponding family-level taxon (or higher rank if the cladogram of a deep level phylogeny is being constructed) prior to the transcription of the figures to the Newick trees, and suprafamilial clade names were also assigned to their corresponding nodes.

During the transcription of phylogenies to Newick notation, poorly supported nodes were collapsed to polytomies when support values are below commonly employed thresholds (70 for bootstrap, 0.95 for Bayesian posterior probabilities, and 3 for Bremer support [Siddall 2002]).

#### Ranking phylogenies

The presence of multiple phylogenetic hypotheses for a given set of taxa seems to be the rule rather than the exception in the scientific literature, especially for biologically well studied taxa. Different phylogenetic trees are found either in different published sources, or even within the same study, in which more than one approach is often used to produce a phylogeny. Considering that such hypotheses are not equally accurate, we assigned a set of criteria for ranking phylogenies for ultimately displaying what we consider the best available tree. Such criteria are based on whether the tree was reconstructed through a cladistic analysis or not, the underlying data for tree reconstruction (type and number of characters), and the inference method, as summarized in Table 1 and as described below.

**Table 1.** Ranking system for selection of phylogenetic hypotheses. The numerical arrangement of each attribute reflects the priorities employed for the selection, starting from the overall approach for the depiction of the phylogeny (in which cladistic character-based analysis are preferred over tentative phylogenies and taxonomic hierarchies), to the last step consisting in the tree inference methodology.

| 1. OVERALL APPROACH | 2. CHARACTER TYPE |  | 3. DATA QUANTITY | 4. TREE INFERENCE METHOD |  |
| --- | --- | --- | --- | --- | --- |
| 1.1. Cladistic (character-based) analysis | 2.1. Molecular | 2.1.1. Nucleotides | 3.1. Number of characters | 4.1. Coalescent approaches |  |
|  |  | 2.1.2. Aminoacids |  | 4.2. Concatenation approaches | 4.2.1. Model based approaches (Maximum likelihood & Bayesian inference) |
|  | 2.2 Morphological | 2.2.1. Molecular scaffold | 3.2. Number of taxa (tiebreaker when number of characters are equal) |  | 4.2.2. Maximum parsimony and distance methods |
|  |  | 2.2.2. Unconstrained |  |  |  |
| 1.2. Tentative phylogeny |  |  |  |  |  |
| 1.3. Taxonomy |  |  |  |  |  |

First, we prioritize trees produced through any character-based cladistic analysis, followed by tentative cladograms proposed by authors. In the absence of any published phylogeny for a given group, taxonomy is then used as a proxy for phylogeny.

Second, character-based cladistic analysis are ranked according to the type of characters employed, in which molecular characters are prioritized over morphological ones given their practical and methodological advantages (Scotland et al. 2003; Vogt et a. 2010). Within molecular analyses, we favor nucleotide characters over aminoacids due to the usually greater informativeness of the former (Townsend et al. 2008). Within morphological phylogenetic analyses, we favor those implementing molecular scaffolds (Lee & Palci 2015) over unconstrained ones.

Third, phylogenies are prioritized based on the number of characters employed for their reconstruction, thus favoring the recent phylogenomic analyses over traditional single-locus or multi-locus approaches. Although it has been shown that a higher amount of characters does not always translate into a more accurate species tree (e.g., Zhang et al. 2021) and are still prone to different sources of systematic error (Philippe et al. 2017; Reddy et al. 2017), we believe that the advantages of larger datasets outweight their drawbacks in most cases, especially considering that recent phylogenomic studies are thoroughly analyzing their data with scrutiny to account for potential misleading issues (e.g., Arcila et al. 2017; Betacur-R et al. 2019; Redmong & McLysaght 2021; Zhang et al. 2021). Higher taxon sampling is used as a tiebreaker in instances of equal number of characters, due to its positive impact in breaking long branches and improving model selection (Nabhan & Sarkar 2012).

Fourth, phylogenies are ranked according to the methodological approach for tree inference, in which coalescent-based approaches are favored over concatenation approaches, due to the former accounting for variation in gene histories by confounding processes such as incomplete lineage sorting (Kubatko & Degnan 2007; Rock & Steel 2015; Jiang et al. 2020). Within concatenation approaches, model-based methods (Bayesian inference and maximum likelihood) are prioritized (Gadagkar & Kumar 2005; Philippe et al. 2005; Puttick et al. 2019), followed by maximum parsimony and distance methods (neighbor joining, minimum evolution). Additionally, analyses that account for further artefacts or systematic errors detected in previous reconstructions are also favored (e.g., compositional heterogeneity across sites in Puttick et al. 2018; long branch attraction in Puttick et al. 2014).

#### Placement of unsampled taxa

Complete taxon sampling is seldom achieved for a single reconstructed phylogeny, which is especially the case for analyses encompassing high amounts of characters such as phylogenomic ones. Consequently, the exclusion of several taxa would be mostly prevalent in the higher ranked phylogenies according to the criteria for choosing a phylogenetic hypothesis. To tackle this issue, we looked for the lower-ranked phylogenies which included each of the missing taxa in the highest-ranked one to infer their phylogenetic placement and incorporate them in the transcribed Newick trees. This step is not straightforward since the topology of such phylogenies may differ from the higher ranked one. In the simplest scenario, to assign the placement of a missing taxon we first identified its sister clade in the lower-ranked phylogeny. Then we looked for such clade in the higher-ranked phylogeny and situated the missing taxon as its sister in the Newick tree (Figure 2a). More complex scenarios arise when the sister clade of such a taxon is absent (i.e., its constituent taxa does not conform a monophyletic group) in the higher-ranked phylogeny. In such instances, we first tagged the constituent OTUs of such a sister clade in the higher-ranked phylogeny, and then we mapped the node representing their most recent common ancestor. The missing taxon is then placed as a direct descendant of such node, resulting in a polytomy (Figure 2b). This is largely the case for assessing the phylogenetic position of fossil taxa, which are mostly derived from morphological phylogenies whose topologies are sometimes drastically different from molecular ones.

**Figure 2.**
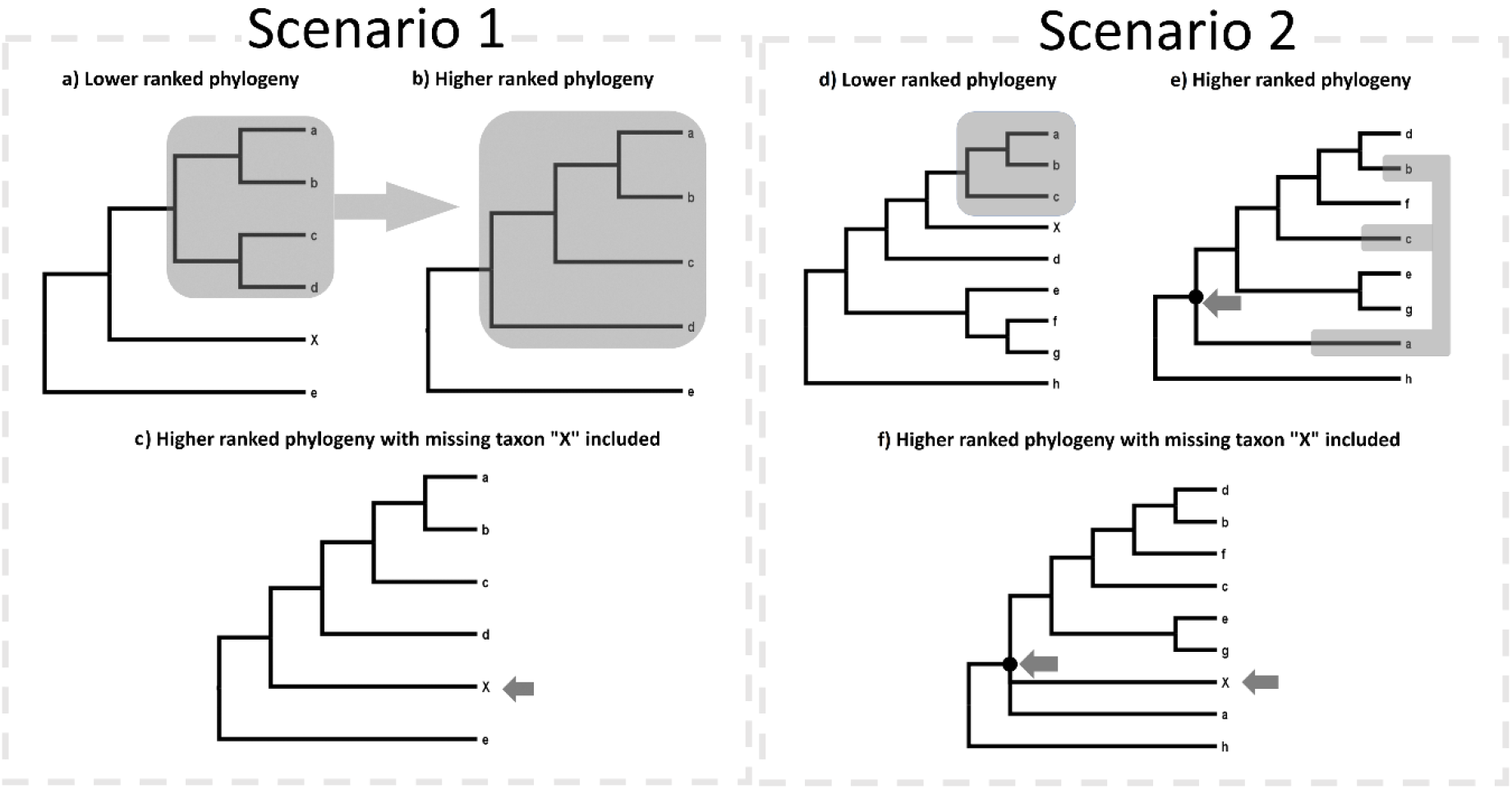
Approach for placing missing taxa in higher ranking topologies. The first scenario consists in (a) a lower-ranking tree which includes the missing taxon X and its sister clade enclosed in gray, and (b) a higher-ranking tree (without taxon X) with this same sister clade also enclosed in gray. In the resulting tree (c), the missing taxon X is then added to the higher ranked topology as sister to the gray enclosed clade. Note that the gray clade is in (a) and (b) is the same in terms of its circumscription (includes *a*, *b*, *c* and *d*), although its internal topology is different. The second scenario also consists of (d) a lower-ranking tree which includes the missing taxon X and its sister clade enclosed in gray, but in the (e) higher ranking phylogeny, this clade does not exist, as its members (*a*, *b* and *c*) are distantly related, conforming a polyphyletic group. In these instances, the node representing the most recent common ancestor between members of the gray-enclosed clade in the lower tree is spotted in the higher tree (gray arrow). Then, the missing taxon X is placed as a direct descendant of such node in the final topology (f), resulting in a polytomy.

#### Non-monophyletic families

One of the consequences of synthesizing supra-specific level phylogenies is dealing with non-monophyletic terminals. This is not uncommon, since for some groups of organisms’ taxonomic classifications have not yet been amended to reflect current knowledge on phylogeny. Non-monophyletic terminals (families in this case) are problematic due to the difficulty of representing accurate topological relationships. Here, we approach this issue with what we consider to be the most conservative solution, as illustrated in Figure 3. For paraphyletic families in which other families of clades are nested within them, we simply allocate such family as sister to their nested lineages (Figure 3a). While the resulting topology would be indistinguishable from one in which all terminals are monophyletic, the paraphyletic status of the family is denoted in the *Remarks* field of the taxon metadata window (see Taxon Metadata for further details)

**Figure 3.**
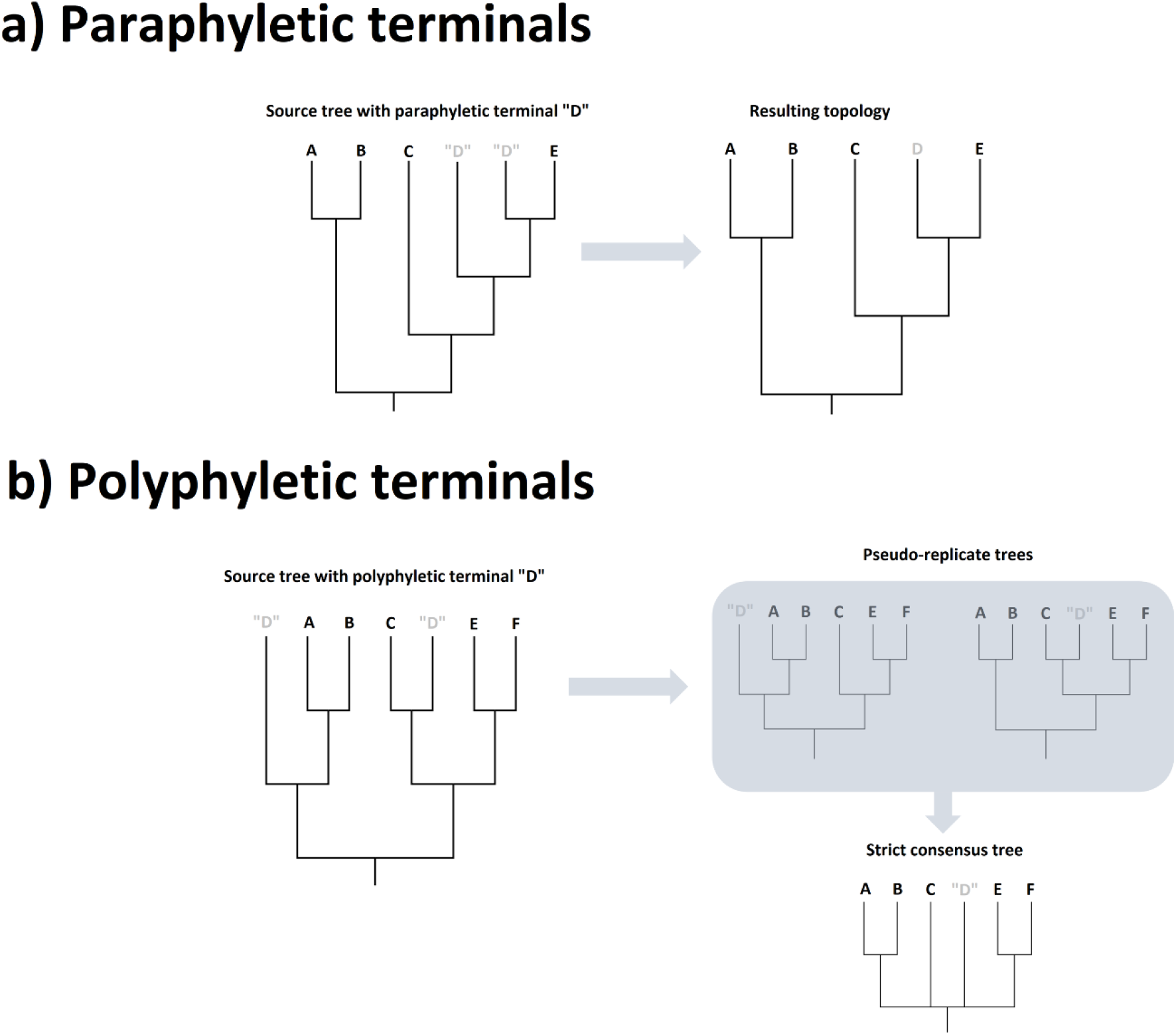
Topological placement of non-monophyletic terminals. a) Source phylogeny with paraphyletic terminal “D”, which in the resulting topology is simply accommodated as the sister lineage of the terminal nested within its grade, which in this case is the E terminal. b) Source phylogeny with polyphyletic terminal “D”. Two pseudo-replicate trees are generated, each with an isolated “D” lineage in one of its two topological positions. A strict consensus is applied for the pseudo-replicates, resulting in a final topology in which “D” is placed in a polytomy. Note the collapse of other nodes in the resulting consensus trees due to the distant position of the “D” lineages”.

For polyphyletic families whose members are distantly related, we pseudo-replicated the source phylogeny N times, where N is the number of distantly related clades belonging to the family. In each pseudo-replicate, only one of such clades is represented, while the others are pruned from the tree. A strict consensus is then applied between all pseudo-replicates (Figure 3b). It is worth noting that this approach may lead to highly polytomic trees when lineages of a polyphyletic family are very distantly located from each other, even when the original phylogeny is highly resolved. This is the case for a few parts of our current synthesized tree such as Microsporidia, in which the highly polyphyletic family Thelohaniidae “breaks up” some of the resolved relationships among other families. This highlights the urgent need to carry out further systematic studies with broader taxon sampling to establish proper taxonomic arrangements. As with paraphyletic families, such instances of polyphyly are also denoted in the taxon metadata text file (see corresponding section below).

#### Time calibration

Dating the entire tree of life poses as an even more difficult challenge than the assemblage of a solely topological tree. The relative scarcity of published time-calibrated phylogenies is the main reason behind such difficulty, leading to a patchy distribution of divergence dates and clade ages estimates across the tree of life. Moreover, the high degree of variation in age estimates across analyses and studies for a given clade also makes this task even more difficult; there are instances in which the age estimate for a clade in a study may be older than the age estimates of its parent clades in another study, which would result in negative branch lengths if assembled in a single tree. This is a serious problem when attempting to synthesize a unified time-calibrated tree of life by compiling estimates from different sources.

To partially overcome such difficulties, we assigned branch length values to our synthesized tree by following two main steps: 1) compiling node age estimates for all possible clades present in such tree by consulting the TimeTree database (Hedges et al. 2009; Hedges et al. 2015; Kumar et al. 2017), and 2) estimating dates of the remaining nodes by using TimeTree data as secondary calibrations, incorporating taxon age ranges from the fossil record and DNA sequences.

We find the TimeTree database to be a suitable source for extracting node age estimates due to (*i*) its extensive collection of data from the literature under properly selective criteria, (*ii*) the mean values reported for divergence dates between different sources, and (*iii*) the consistency tests performed to avoid negative branch lengths. Data collection was manually performed between June 2017 and December 2019 by querying taxon names, as the whole database is not available for download. Despite the strict selection criteria employed by the authors to assemble the database, we nonetheless filtered out node ages that were younger than the oldest occurrence of the current fossil record.

To estimate branch lengths and divergence dates of the remaining nodes not covered in the TimeTree database, we compiled age range estimates from the fossil record for each family by consulting the PBDB between 2014 and 2021. Age ranges obtained from alternate sources were cited for each taxon in the taxon metadata text file (see next Methods subsection) for their eventual display in secondary information windows. We carefully excluded some spurious records for taxa whose ages dramatically expanded their ranges (i.e., were too old or too young) by revising the recent literature to address whether those records represent dubious taxonomic assignations. The resulting age ranges for each taxon were then used to filter out some of the node ages compiled in the previous step.

Nucleotide sequences were also obtained from GenBank for every family in the synthetic tree present in the database. Because there is little overlap of homologous loci between different major groups across the tree of life, we partitioned the tree in 20 subsections, with some of them nested within others. For each partition, we selected the set of loci that maximizes taxon coverage, so for some partitions more than one locus was necessary (Table 2). The procedures described hereafter were performed separately for each tree partition.

**Table 2.** Tree partitions and their respective loci employed for the phylogeny-dating analysis in BEAST2. Nested partitions are indicated within parentheses. Accession numbers for family-level taxa are listed in Supplementary Table 2. *Metazoa and Hexapoda have nested clades that were treated as separate partitions (Porifera and Endopterygota, respectively) despite the use of the same set loci. This was necessary due to their high number of terminal taxa and the consequent difficulty in running the analysis with large data matrices.

| Tree partition | Loci |
| --- | --- |
| Prokaryotes (Bacteria and the "archaeal" grade) | 16s |
| Eukaryota (excluding Chlorophyta, Streptophyta, Amoebozoa, Fungi and Metazoa) | 18s |
| Metazoa* (excluding Porifera, Xenocarida, Hexapoda and Vertebrata) | 18s |
| Porifera | 18s |
| Xenocarida | COI, H3 |
| Hexapoda* (excluding Endopterygota) | COI, 18s |
| Endopterygota | COI, 18s |
| Vertebrata (excluding Chondrichthyes, Actinopterygii, Squamata and Passeriformes) | Cytb |
| Chondrichthyes | Cytb, NADH2 |
| Actinopterygii (excluding Characiphysi) | COI |
| Characiphysi | Cytb, COI, 12s |
| Squamata | Cytb, 16s |
| Passeriformes | Cytb, COI, RAG1, Myo |
| Chlorophyta | rbcl, 18s |
| Streptophyta (excluding Bryophyta and Malvales) | rbcl |
| Bryophyta | rbcl, nad5 |
| Malvales | rbcl, 18s |
| Fungi (excluding Ascomycota and Chytridiomycota) | 18s, 28s |
| Ascomycota | 18s, 28s |
| Amoebozoa | 18s, Actin |

**Abbreviations:** 12S=12S ribosomal RNA, 16S= 16S ribosomal RNA, 18S= 18S ribosomal RNA, 28S=28S ribosomal RNA, COI= cytochrome oxidase subunit I, Cytb=cytochrome b, H3= Histone 3, Myo=myoglobin, NADH2=NADH dehydrogenase subunit 2, nad5=NADH dehydrogenase subunit 5, RAG1=recombination activating protein 1, rbcl=Rubisco large subunit,

Sequences were aligned and concatenated (when presenting more than one locus) with Geneious^®^ v5.4 (Drummond et al., 2011) using the MUSCLE algorithm (Edgar 2004), and conflicts were manually resolved by visually inspecting the resulting alignments. For a total evidence dating analysis of tree partitions, alignments were imported to the utility program BEAUTi for setting the parameters of BEAST2 (Bouckaert et al. 2014) utilizing a fossilized birth-death model (Heath et al. 2014). To add taxa with reported age ranges from the fossil record and no molecular data, we edited the resulting alignments for manually adding such taxon names with empty sequences. Ages for fossil taxa were set (in millions of years) as the younger bound values of their ranges, and an uncorrelated lognormal relaxed clock was used as the clock model. Monophyly constrains were applied to match the topology to that of the synthesized tree, and the compiled node ages were then applied as secondary calibrations, setting them as mean values under a normal distribution with a standard deviation of 0.0001, to constrain the node ages of the resulting tree to the original TimeTree values as much as possible.

Because models that account for stratigraphic ranges (rather than uncertainty ranges) have not yet been implemented for phylogenetic analysis, we could not directly establish the older bounds of familial age ranges. Consequently, to incorporate these older bounds in an indirect way, we used uniform calibration priors on nodes whose direct descendants include at least one taxon presenting age range information. The minimum bound of the distribution was set as the older bound of the taxon’s age range, and the maximum bound was set as the age of the nearest normally distributed, secondarily calibrated parent node (Figure 4). If more than one of the descendants of a node had age range data, the minimum bound of the uniform distribution was set as the oldest value of such age ranges.

**Figure 4.**
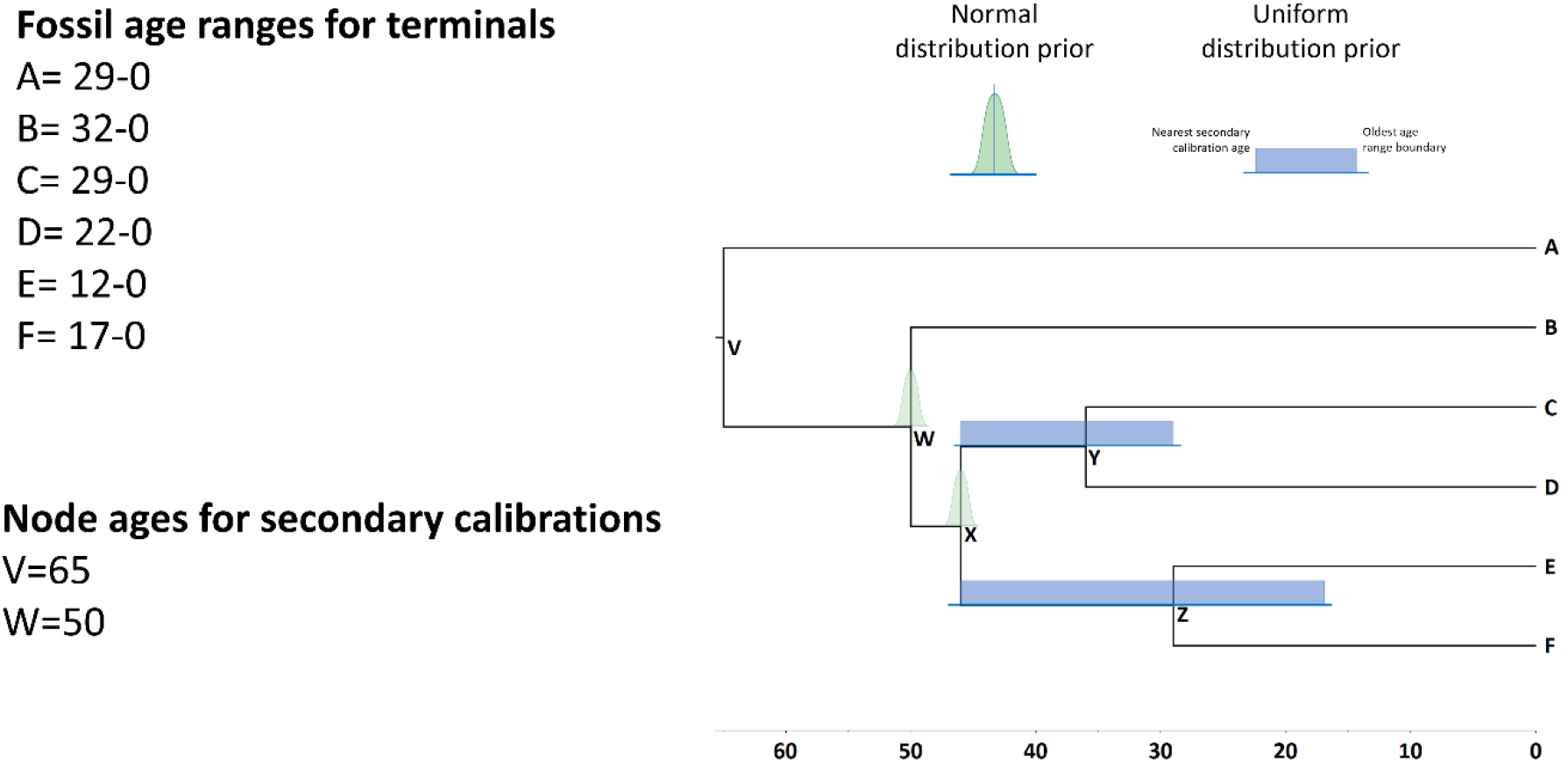
General approach for dating the phylogeny in BEAST2. In this hypothetical phylogeny, all six extant terminals exhibit sequence data and age ranges from the fossil record, spanning from the present (0) to their oldest fossil occurrence in the past. Additionally, two of all five nodes have reported age estimates from the literature (V and W). These estimates are employed as secondary calibration points, setting them as the mean value of normal distribution priors (with very narrow standard deviations). Additional uniform distribution priors are set for the nodes whose direct descendants are terminal taxa with reported age ranges (Y and Z). The lower bound of the uniform distribution is set as the oldest bound of the ranges of its daughter terminal taxa: (29 for node Y and 17 for node Z, from C and F terminals, respectively). The upper bound of the uniform distribution is set as the nearest secondarily calibrated parent node, which in this case is W (age=65) for both cases. Note that in this hypothetical scenario, age ranges for taxa A and B are not utilized because they are direct descendants of secondarily calibrated nodes (and whose maximum age values do not exceed such secondary calibration ages anyways). Moreover, age ranges from taxon D and E are not employed either because their respective sister taxa have older age values in their ranges, which are ultimately employed for the uniform distribution priors.

Because of the constant need to re-run these analyses each time the synthesized tree is updated, we elected a GTR+I+G substitution model for all analyses (to avoid time expenditure by testing for multiple models), as it is one of the models that most frequently fits real data (Arenas et al. 2015). Each analysis was run for 1.000.000 generations, sampling every 1000 trees. The maximum credibility tree was generated using mean values with the program TreeAnnotator in the BEAST package, with a 10% burn-in cutoff. The resulting time-calibrated trees were then used to manually assign branch lengths to the synthesized Newick tree, rounding values up or down to the closest integer.

Despite all the collected data to obtain age estimates for every node in the synthesized tree, there are several family-level taxa with neither molecular data nor stratigraphic age ranges from the fossil record. These taxa were excluded from the previously described procedures, and the age of their parent nodes were manually calculated by equally dividing the distance between the nearest parent and daughter dated nodes, which is an identical approach to the one employed in the BladJ algorithm (Webb et al. 2002). Branch lengths were then assigned in accordance with these interpolated node ages.

### Taxon metadata

For each of the families and supra-familial taxa in the resulting synthetic tree, we gathered information on their common names and taxonomic synonyms. Most common names were taken from the NCBI taxonomy (Federhen 2012), Catalogue of Life (2021), Encyclopedia of Life (Parr et al. 2014), and Christenhusz et al. (2017; specifically for vascular plants). Synonyms were taken from the NCBI taxonomy, PBDB, World Register of Marine Species, Wilson & Reeder (2005; for mammals), Stevens (2016; for vascular plants), and Kluge (2010; for supra-familial insect taxa). Supporting sources for phylogeny and taxonomy were also annotated for each taxon, along with the non-monophyletic status of some families. Taxon metadata was stored into two separate text files, one for common names, and another for the rest of the attributes that are eventually displayed in primary (common names) and secondary (sources, synonyms, and others) information windows (see Mobile App Overview section in Results).

### Selection of pre-collapsed clades

In order to exhibit a backbone of the tree of life to easily illustrate the relationships between major groups of organisms, we decided to collapse a set of higher-level taxa so the initial view is rendered with 88 terminals. These higher-level taxa were not necessarily at the same taxonomic rank, but where rather chosen arbitrarily to match what we consider some large and popularly known groups or organisms, and whose names were listed in a plain text file as one of the input files for the tree rendering code (see this taxon list in Supplementary File 1).

### Taxonomic coverage

The methodological procedures described up to this point were applied over 68 of the 88 terminals shown in the initial backbone tree, as specified in the previous section (Selection of pre-collapsed clades). The 20 clades that have not been covered yet are listed in a plain text file (see list in Supplementary File 2) that is used as an input for the code of the application, whose purpose is to denote its current unavailability to users. These taxa also appear as pre-collapsed clades, but unlike others, are unable to be uncollapsed by the Open button (see Mobile App Overview section in Results).

### Illustrations

Family-level images were produced by eight different artists (listed in the Credits section of the mobile app) and were loosely based on a set of real-life photographs and/or artwork material provided by the authors of this study. Due to the high number of images needed and the limited storage of mobile devices, we opted for rather simple but faithful artistic representations of the organisms, instead of overly detailed illustrations. Each illustration was based on a representative member of the family, which was often a species of its type genus. Because of the lack of specific paleontological knowledge of the artists, illustrations of fossil taxa were always based on pre-existing artistic reconstructions. Consequently, we excluded taxa based on very fragmentary remains that have not been artistically reconstructed to the date of this publication.

When possible, we attempted to depict the same sex and life stage within groups of closely related taxa. For instances of marked sexual dimorphism, we chose the sex that mostly retains the general body-plan of closely related taxa. Using the case of coccoid insects (Hemiptera: Coccoidea) as an example, in which females are wingless and highly paedomorphic individuals, we chose the winged males for the easier visual comparison of homologous traits with other hemipteran families and insects in general. We also chose adult stages for plants and animals, the life stage that produces fruiting bodies for multicellular fungi, and trophozoites for apicomplexans. Finally, although it is common practice for illustrations of modular organisms such as plants to depict focused parts of the individual, we instead attempted to depict the entire organism for those presenting large, woody growth habits such as trees, shrubs, and hedges.

Images for supra-familial clades were generated by transforming an image of one of its daughter families into a gray silhouette. Each representative family was chosen based on what the authors consider to be the most ancestral morphology for the whole group, as very specialized morphologies may not be a good representation of an entire clade.

### App Development

A mobile app for interactively exploring and visualizing the Newick tree was written in Dart with Flutter 2.2 (https://flutter.dev/), an open-source software development kit employed to program cross platform applications (incluging Android and iOS). The code for tree rendering was loosely based on that of Figtree v1.3.1 (Rambaut 2012). The resulting code employs a set of text files as input, which include the Newick tree file, the Common names file, the Taxon Metadata File, the Pre-Collapsed Groups file, and the Unavailable groups file, as well as an image folder containing the whole set of taxon icons in Portable Network Graphics (PNG) format.

## RESULTS

As of May of 2021, the assembled tree comprises 7335 terminal taxa (Figure 5; for a detailed view, see Supplementary Figure 1), with 7315 of them being families, while the remaining 20 being higher level groups whose family-level phylogenies have not been assembled yet (Annelida, Echinodermata, Brachiopoda, Branchiopoda, Bryozoa, Cnidaria, Chelicerata, Ciliophora, Dinozoa, Endomyxa, Filosa, Heterokontophyta, Multicrustacia, Myriapoda, Oligostraca, Mollusca, Nematoda, Plathelminthes, Retaria and Rhodophyta). Moreover, 1999 suprafamilial clades were included and annotated as node labels in the Newick tree.

**Figure 5.**
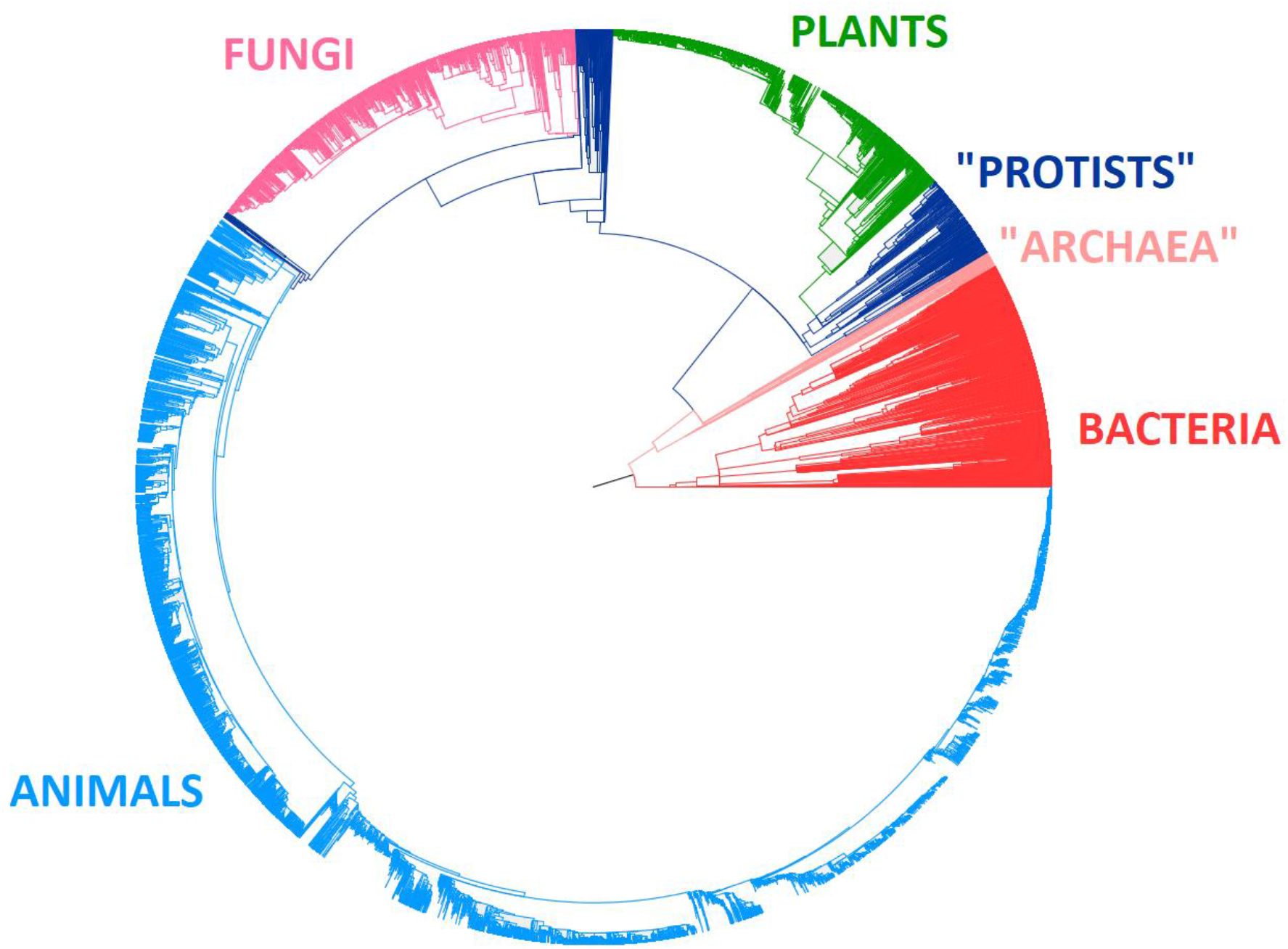
Current version of the family-level synthesized tree with six major groups of organisms coded by color, with non-monophyletic groups enclosed in quotation marks. Branches that do not reach the outer rim of the circular phylogeny represent fossil families, which are especially evident in animals and absent in bacteria, archaeans and protists.

At the time of publication, a total of 1479 bibliographic sources have been employed to assemble the final time-calibrated tree (see Reference section in the mobile app or Supplementary File 3), excluding those that were revised but eventually discarded because of the availability of higher ranked phylogenies. The time calibrated tree comprises 559 secondarily calibrated nodes obtained from the TimeTree database (Supplementary File 4), while age ranges from the fossil record were acquired for 2859 taxa (Supplementary File 5) and molecular data for at least one locus were obtained for 5060 taxa among all tree partitions (Supplementary File 6). Neither molecular data nor stratigraphic age ranges from the fossil record were available for 333 families. Of the 4888 nodes in the resulting synthetic tree, the age of 4084 of them were estimated through the BEAST2 analyses, while the remaining ones were fixed as secondarily calibrated nodes (from TimeTree) and nodes whose age values were assigned by interpolation.

Regarding taxon metadata, common names were obtained for 2389 taxa, and 797 taxa presented at least one taxonomic synonym. Moreover, 126 families were found to be non-monophyletic as currently circumscribed according to the revised source studies.

In addition to the code for tree rendering and its interactivity features, the resulting APK file of the developed mobile application includes the synthesized Newick tree file, the common names text file, the additional taxon metadata text file (phylogenetic and taxonomic sources, synonyms, and remarks), the pre-collapsed clade list text file, the unavailable clade list text file, and the 9334 illustrated icons for both families and supra familial clades, totaling up to 128 megabytes. The app is freely available for download in Android devices in https://play.google.com/store/apps/details?id=com.pappcorn.tol and in iOS devices in https://apps.apple.com/co/app/tree-of-life-tol-app/id821256110

### Mobile App Overview

The main menu of Tree of Life App offers five main options: Explore, Tutorial, References, Frequently Asked Questions (F.A.Q.), and Credits (which lists all the staff who participated in the project). Most of the following contents is focused in the Explore option, which revolves around the exploration of the large phylogeny assembled here.

The Explore section (Figure 6) leads users to a circular phylogeny representing the family-level tree of life. Exploration is performed by the standard zooming and panning using the standard gestures on touch screens of mobile devices. Instead of exposing all the family level content, the initial view of the tree consists in several large clades of organisms displayed as collapsed branches, so users can easily visualize the higher-level relationships across the tree of life before straying in an overwhelmingly large tree diagram (Figure 6a). These pre-collapsed branches do not represent any specific taxonomic rank but were rather arbitrarily selected to represent large and well-known groups. Users may either select these collapsed branches individually to unfold the desired family-level relationships or employ the “open all” button at the upper right of the screen to unfold the entire family-level tree (Figure 6b and c).

**Figure 6.**
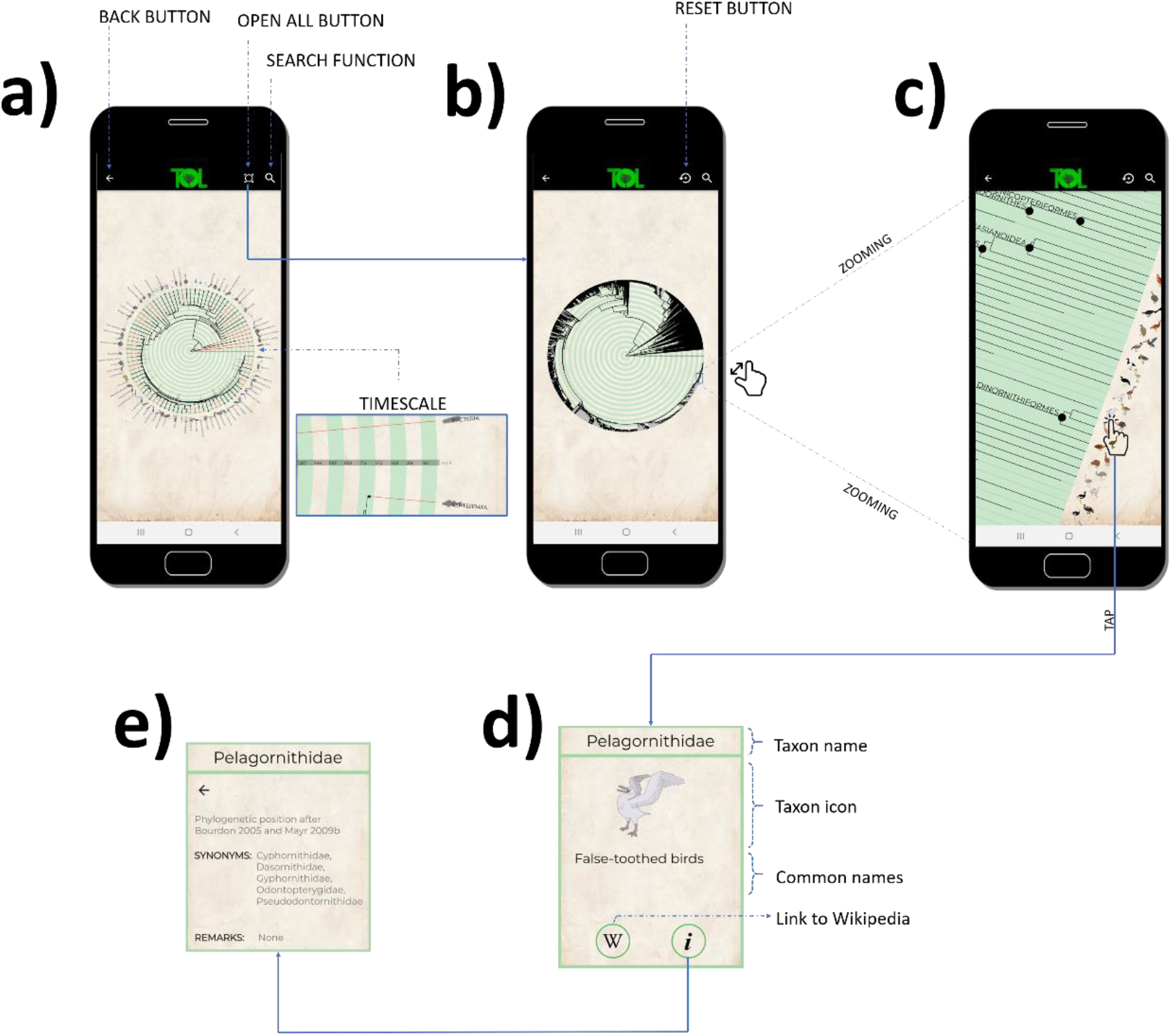
Basic features of Tree of Life app. a) Initial view of the tree with major groups of organisms pre-collapsed into single branches. b) A full view of the family-level tree of life after activating the “open all” button. c) A zoomed in portion of the tree focused on part of the paleognath and galloanseran birds. c) Primary information window displayed after tapping on one of the pictures at the tips. b) Secondary information window after tapping on the “*i*” button.

In contrast to other efforts to compile the tree of life in which only topological information is shown, we provide a time-calibrated phylogeny that might be used to visualize divergence dates among lineages, as well as extinction dates of fossil taxa. This is achieved with the aid of the radial ring pattern on the background and the horizontal labeled timescale at the right hemisphere of the circumference (Figure 6a).

Furthermore, unlike previous tools, Tree of Life App does not depend on internet connection for its main functionalities as all data is stored within the device once the app is downloaded. Consequently, users might explore and consult the tree of life in remote locations (i.e., during field work) or situations in which internet connection is scarce or lacking.

#### Branch tips

One of the most distinctive features of the phylogeny displayed in Tree of Life App is that rather than plain text, illustrated icons of organisms are displayed at the branch tips of the tree. Displaying images instead of text can be advantageous for several reasons, not only because they make the tree more visually appealing, but also because an image of the organism is arguably more informative than the taxon name when the latter is mostly unknown for an average user. Consequently, we argue that an image-rich tree of life is the best way of visually exhibiting the evolutionary history of life, which is in line with the recent ongoing trend of displaying tip-illustrated phylogenies in the figures of published studies (e.g., Hasegawa 2017).

Two types of icons can be seen at branch tips: color icons and gray silhouettes. Color icons correspond to taxonomic families, which are OTUs at which this tree was constructed. The organism displayed in an icon correspond to a representative species of each taxonomic family. As one of the several interactive features of this app, these icons can be tapped on to access what we call a primary information window (Figure 6d), which displays the enlarged icon, the taxon name, and its associated common names, if any. Furthermore, a link to Wikipedia is also present at the lower right corner of the window for users that intend to get further information of the selected taxon, as well as a button that leads to a secondary information window (Figure 6e) containing the sources supporting the phylogenetic position of such taxon as well as other bits of information (see further details in the References section of this article). The second type of icons are gray silhouettes, which in contrast to the colored ones, have their corresponding taxon name placed over the image. Rather than taxonomic families, these icons correspond to supra-familial taxa that are either the pre-collapsed groups shown in the initial view of the tree, or manually collapsed clades obtained by using the branch options described in the next section. Users may also tap on these silhouette icons, but the options shown in their corresponding pop-up windows differ from those of colored icons as described below.

#### Internal Node Options

In addition to the family-level branch tips, internal nodes are also labelled on the phylogeny denoting supra-familial taxa or clade names. Such labelled nodes are displayed as black dots adjacent to their text label (Figure 7a). Users may tap on the black dots to display a primary information window which, as in the case of their branch tip family-level homologues, display a gray silhouette icon denoting a representative organism of the clade, its associated taxon name, a link to Wikipedia and the References section, and two additional options exclusive to supra-familial taxa: Subtree and Collapse (Figure 7b).

**Figure 7.**
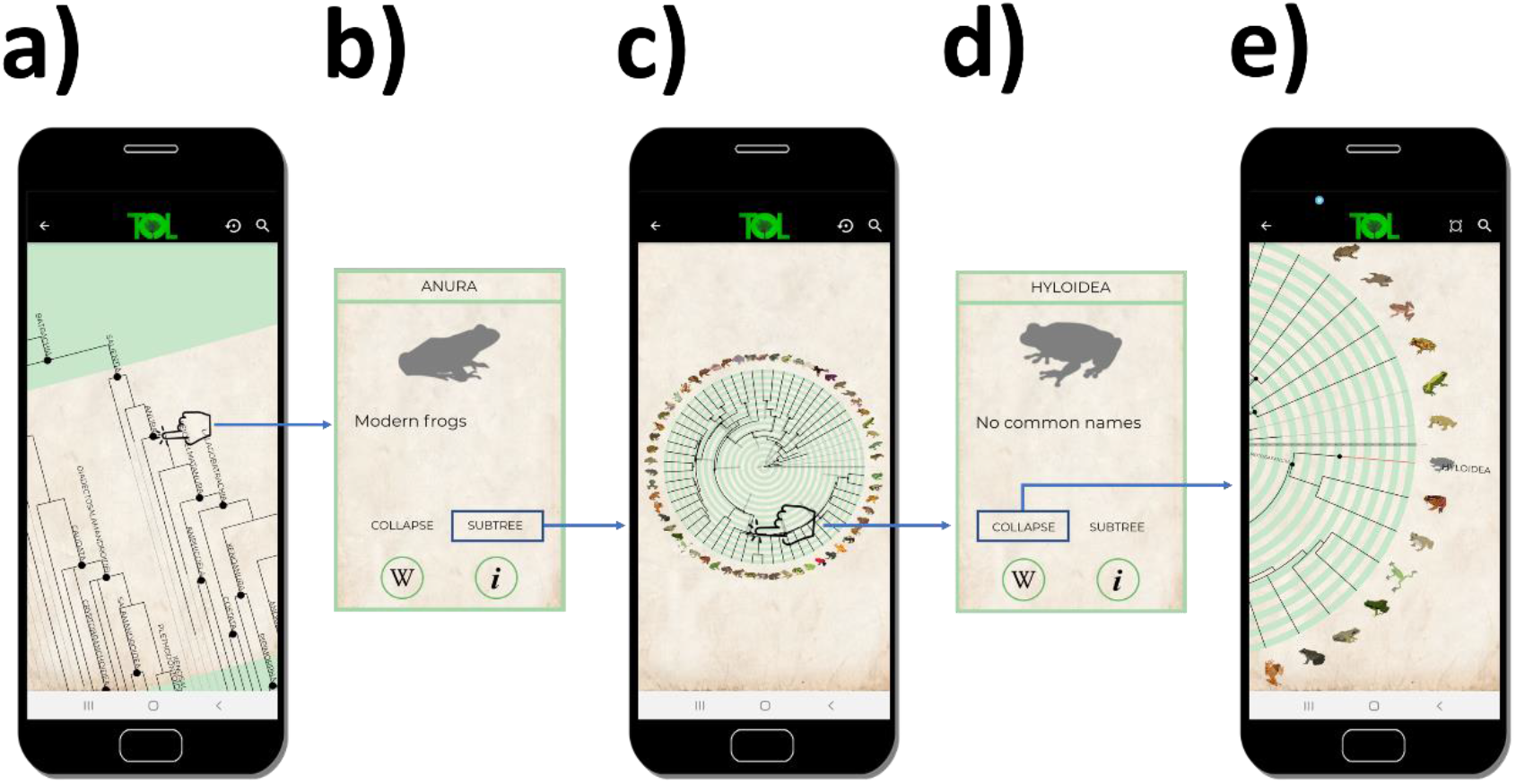
Some examples of branch options. a) A zoomed in portion of the tree focused on the deep nodes around anurans (frogs). b) Primary information window displayed after tapping on the Anura node, displaying both branch options at the bottom of the window. c) Full view of the anuran tree after selecting the Subtree option. d) Another view of a primary information window, this time selecting the Hyloidea node within the anuran tree. e) Appearance of the anuran tree after selecting the “Collapse” option on the Hyloidea node. Note that Hyloidea is now represented by a single red branch with a gray silhouette and text label.

##### Subtree

Users may employ this option to visualize specific clades of the tree by excluding everything outside the selected clade. In such cases, smaller circular trees may be displayed without the need of visualizing them by thoroughly zooming in a specific location of the large tree of life (Figure 7b,c).

##### Collapse

This option enables the user to collapse the entire clade denoted by the selected node into a single branch. The collapsed clade or branch will then be displayed as a red line with a gray silhouette at its end, along with a text label displaying its taxon name over the icon (Figure 7d,e). This feature allows the user to customize the tree at will by hiding undesired details of the tree for any specific purpose, (e.g., visualizing the tree at higher, supra-familial levels by selectively collapsing different clades). To re-open a collapsed branch, the silhouette icon at the tip must be tapped, and then an almost identical pop-up window will appear, with an “open” button replacing the previous “collapse” button must be tapped on to unfold the contents of the collapsed clade.

##### Highlighting Branches

Highlighting single branches or even entire clades is possible by tapping pictures at the tip or black dots at the nodes and then closing the primary information window. The selected branch will then appear highlighted in blue. This may be useful to keep a lineage marked without losing sight of it in a highly dense tree, to easily identify the sister group of a lineage by tracking the selected branch to its parent node, or to view the family-level diversity of a clade relative to the rest of the displayed tree.

#### Secondary information windows

Users may access secondary information windows by tapping on the “i” buttons in each of the primary pop-up windows that are accessed by tapping on icons at the branch tips or labelled nodes. These windows contain three information fields: a first one for the citations of sources supporting the taxonomy, phylogeny, or phylogenetic position of the taxon, a second one for taxonomic synonyms, and a third one for additional remarks such as the phyletic status of a family (Figure 6b).

#### Search Function

As the family-level tree of life displayed is considerably large, users may have a hard time finding specific organisms in such a big tree. To solve this problem, a search function is available so users may query specific taxa by typing either taxon (both family names and higher, supra-familial clade names) or common names. A green arrow tip will then appear indicating the location of the matched results. In the current version of the app, the search function is unable to find queried taxa if they are within a collapsed group. Given that the initial view of the tree of life consists in large pre-collapsed groups, this limitation implies that users must have an *a priori* knowledge regarding the higher-level taxonomic affiliation of the queried taxon, and then open its corresponding branch before using the search function. However, users may use the “open all” button (Figure 6a) to unfold all the pre-collapsed branches shown in the initial view to visualize the whole family-level tree. This way, the search function will successfully find any taxon included in the current version of the phylogeny.

#### References Section

The tree of life shown in the app was synthetized from over 1400 sources that provided phylogenetic and/or taxonomic information. Each of these sources are properly cited in the secondary information windows and listed in the References section of the main menu, which are arranged alphabetically and divided into six separate pages with their respective alphabetical ranges.

## DISCUSSION

By a thorough revision of the scientific literature, a set of rigorous criteria for source selection, and a collection of divergence dates, fossil stratigraphic records and sequence data, we assembled an up-to-date family-level tree of life. Additionally, we developed an app for mobile devices to interactively explore and visualize the large, synthesized tree. To our knowledge, this is the only project that have approached data compilation and visualization issues at the same time, as other similar initiatives have relied on others for one or the other (LifeMap and OneZoom with the NCBI taxonomy and the OTOL synthetic tree). In the following subsections, we discuss the strengths, novelties, limitations and pitfalls of the performance, interactivity features, and phylogenetic information exhibited in this project.

### Strengths and novelties

#### Fossil taxa

A relevant novelty introduced in our synthesized tree is the inclusion of fossil taxa, considering that large-scale phylogenetic data sources employed by other tree of life visualizers (the OTOL by LifeMap and OneZoom) are currently devoid of extinct organisms. The inclusion of fossil taxa into phylogenies have long been considered crucial for the understanding of overall evolutionary processes (Prothero 2007; Lyson et al. 2010; Slater et al. 2012). By breaking long branches, the incorporation of fossil taxa provides a clearer visualization of the gradual evolution of traits because of the transitional morphology of some extinct organisms. This is further exploited in our mobile app by the representation of branch terminals with images. In this way, users can easily observe gradual transitions between disparate morphologies, as well as understanding the close relationships between some extant taxa that would otherwise be surprising (e.g., birds to other non-avian “reptiles”, cetaceans to hippopotami, terrestrial vertebrate to fish).

#### Branch lengths

Another novelty strongly tied to the inclusion of fossil taxa is the implementation of a temporal axis and time-proportional branch lengths. Visually inferring branch lengths and divergence dates provide valuable information that complements topological visual cues; while the latter indicates plain relatedness, the former allows user to visualize the degree of such relatedness. While this is also lacking in the OTOL synthetic tree, it is nevertheless unable to be visualized by the formats displayed by OneZoom and LifeMap. It is worth mentioning that OneZoom provides ages for many of its nodes with numerical labels, but the age of such clades cannot be visually inferred by the proportionality of branch lengths. The TimeTree project also assembled a large time calibrated tree, but such a project is mostly employed as a divergence date database rather than as a topological tree assemblage or for tree exploration purposes, due to their restricted coverage and the narrow range of eligible studies (only time-calibrated phylogenies, which are a small fraction of the overall published phylogenetic knowledge).

#### Images

Partially in accordance with what was previously mentioned about visualizing fossil taxa in phylogenies, we consider that one of the most solid strengths of our version of the tree of life is the representation of terminal taxa in the form of illustrated icons. We argue that this feature not only enhances the aesthetic appeal of the tree, but also provides valuable visual information to users who are not familiarized with certain groups of organisms. This could even prove attractive enough for the non-academic public who may not even be able to properly read phylogenies but might very well be engaged to visually explore the morphological diversity of life arranged by evolutionary relatedness.

The incorporation of images in a phylogeny is not particularly a novelty, as it is not uncommon to find publications with organisms’ illustrations or photographs in their phylogenetic figures. In fact, the OneZoom visualizer incorporates some images within its leaves and internal nodes, consisting mostly in circularly cropped photographs. Although photographs have a huge advantage over illustrations because they faithfully represent the real-life image of an organism, we find this approach as a disadvantage in other aspects. First, photographs are usually presented as a geometrical area (squares, rectangles, or circles in the case of OneZoom) whose contents also include a background behind the depicted organism. This is problematic because the true shape and outline of the organism itself is gradually lost and becomes progressively indistinguishable from the rest of the photograph while zooming out, and zooming out is ironically crucial for a broad comparison of multiple organisms in a phylogeny. Additionally, photographs may include only specific parts of the organisms and some of those may not even be homologous with the parts displayed in other photographs, so a proper comparison is not always achieved. By presenting images without a background —as is the case with the illustrated icons presented here— the shape of the organism is easily maintained over a broad range of zoom levels, enabling its identification and comparison with its neighboring taxa. As our icons also depict the whole organism and mostly homologous life stages, a proper comparison is achieved while visualizing many related organisms at once within the phylogenetic framework.

#### Literature coverage

Incorporating as much of the relevant literature as possible is pivotal for assembling a tree of life that best represents most of the generated knowledge on the subject. However, prevailing large-scale projects rely on specific data formats for the inclusion of studies into their synthetic tree, limiting the number of eligible sources and consequently, representing only a fraction of all the available knowledge on phylogeny. The OTOL is notorious for this restriction because its infrastructure was designed to exclusively incorporate phylogenetic data in the form of tree files, but the overall presence of this data format in publicly available repositories is low, and requests to authors for such files had a low response rate (Hinchcliff et al. 2015). Because of this, the placement of 98% of the tips in their synthetic tree is based on taxonomy rather than phylogenetic studies at the time of its formal publication, although this number may have been reduced since then. On the other hand, the TimeTree project is focused on providing a summary on divergence time estimates and as such, it incorporates a small fraction of the phylogenetic knowledge, which consist in time-calibrated trees. Moreover, it is also dependent on tree files obtained through public repositories or by directly contacting authors, presumably suffering from the same difficulties mentioned for the OTOL project.

Contrastingly, we bypassed these obstacles by manually transcribing phylogenies available as figures to Newick notation, allowing us to incorporate virtually any published study into the tree assembling process. Hence, we argue that the tree of life displayed in the app is a comprehensive representation of the available knowledge on phylogeny in the scientific literature.

#### Tree layout

Rectangular or diagonal tree formats are the most commonly used layouts for publishing phylogenies, and circular formats are often used for large phylogenies. Despite this, current tree of life explorers —namely OneZoom and LifeMap— employ unfamiliar tree layouts to display their extremely large phylogenies. These layouts may facilitate exploration by deep zooming, but the reading of topological relationships is troublesome and the overall shape of the tree at shallow zoom levels is hardly perceived (i.e., the relative size of clades and tree asymmetry).

We attempted to circumvent these pitfalls by representing our synthesized tree in a traditional and conventional format: a circular phylogeny. This layout is useful for optimizing space for large trees and provide the additional advantage of preventing the usual interpretation of phylogenies as evolutionary ladders, rather than a branching process. As with other conventional formats, this layout also allows the incorporation of a linear timescale, which in this case displays time-proportional branch lengths.

#### Internet connection

Lastly, another relevant novelty of our tree of life explorer is its independence of internet connection, which contrasts with other prevailing explorers in their reliance to web servers. The tree file, taxon metadata, reference list and organisms’ icons are all stored within the mobile device once the app is downloaded. The only features that require internet connection are the tutorial (which redirect users to a YouTube video) and the links to Wikipedia for each taxon, which is shown in primary information windows.

Nevertheless, the local storage of all these files (especially images) has a relevant drawback. The independence of a web server combined with the limiting processing power of mobile devices (in contrast to computers) leads to a relative loss of performance for features such as the smoothness of zooming and panning gestures. This performance difference issues are especially evident in older devices when compared with the superior exploration capacities of the OneZoom and LifeMap explorers.

### Limitations

#### Current taxonomic coverage

Although the tree assembling procedures described here allows a broad sampling of the available phylogenetic knowledge, these thorough processes have only been carried out over selected group of organisms due to the time-consuming methodologies. As such, with over 7200 terminal taxa, around half of the total family-level diversity is represented at the moment (Ruggiero 2014; PBDB). Some of the large and well-known groups among those currently unavailable sections of the tree are chelicerates (arachnids and allies), multicrustaceans (the largest clade of the now paraphyletic Crustacea), mollusks, annelids, cnidarians, and red algae, along with other smaller or less popular groups.

Additionally, within the large, covered groups, some taxa are absent because either they have not been formally assigned to a family-level taxon or no photograph, image, or artistic reconstruction exists for such taxa. The former case is especially evident in some fossil taxa whose researchers have largely abandoned Linnean ranks, such as current systematics of sauropod dinosaurs. For representing those groups in our tree, we included some older, family-level described taxa, but newly described taxa from the last two decades are unfortunately unrepresented. This is also the case for some family-unaffiliated taxa denominated as *incertae sedis* (most usually found in microscopic fungi and algae, as well as some unicellular eukaryotes), although these organisms represent a minuscule fraction of the described biodiversity. On the other hand, the lack of imagery is notorious for fossil taxa based on very fragmentary remains, which makes the depiction of the whole organism difficult. This is also the case for candidate taxa that are only known from environmental DNA samples, leading to the exclusion of a vast amount of the known prokaryote diversity in the tree of life (Hug et al. 2016). Although these taxa could be incorporated as branches with only text labels in their tips, we consider this approach would not fit within this project’s scope of visualizing a fully illustrated tree of life. For this reason, we refrain for including such taxa until more complete fossils are discovered, or some of the candidate taxa become cultivated and formally described (e.g., Imachi et al. 2020).

#### Taxonomic resolution

The fact that the goal of this project is to produce an illustrated, family-level tree of life implies that a broad variation of the diversity of life is to some extent reduced to single terminals representing clusters of closely related species. Hence, phylogenetic relationships below family level are out of the scope of this project and should be consulted in other sources encompassing a higher taxonomic resolution, such as the OTOL.

#### Divergence date estimates

Unlike reconstructing cladograms or phylograms from sequence data, dating phylogenies is a more complex and difficult process, as it requires considerable *a priori* knowledge on the detailed fossil record of the specific clade being treated and the tentative systematic position of fossil taxa within the tree. Due to our lack of this specialized expertise, we attempted to use as many secondary calibration points as possible, so several of the node ages in the resulting tree represent estimates from other studies. However, we did attempt to estimate many other nodes in between those secondary calibrations, and although we obtained (and to some extent curated) stratigraphic range data from the PBDB for a wide array of taxa, several inaccuracies from such database might have been overlooked. Moreover, the fact that the BEAST analyses we carried out were run for relatively few generations, along with the lack of chain convergence for some runs and the use of a single substitution model for all tree partitions without prior model testing, implies that some node ages estimated here should be interpreted with caution.

### Future projections

Besides the constant incorporation of novel and better ranked phylogenies published in the literature, our most relevant short-term goal involves covering the currently unavailable groups for displaying a fully comprehensive tree of life.

Several other additional features for the mobile app are also under plans, such as additional taxon associated metadata such as number of described species; the incorporation of a geologic time scale with all its conventional subdivision, parallel to the already implemented numerical timescale; a function to find and highlight the path and the divergence date between two selected taxa; an educational and didactic section for enhancing tree-thinking skills, and a trivia game section in the main menu where a random set of taxa is drawn from the tree and users must establish their proper topological relationships. Finally, a web app is also under consideration so users can access this tool from their laptops and desktop computers.

## ACKNOWLEDGEMENTS

This project could not have been completed without all the talented professionals who were essential for the development of this project, most notably the artists Katie Walker, Vanessa Reina, Oscar Builes, Camilo Delgado, Lina Beltrán, Luisa Orozco and Carolina Salazar, the developers Mario García, Janet Brumbaugh, Alex Lamb, Yang Xiaoming and Nicolas Contreras, and the graphic designer Ricardo Mejía. We also thank the PappCorn S.A.S. team for its overall guidance into the world of mobile app development. Finally, we are very grateful to Adolfo Jara, Maria Sole Calbi, Francisco Fajardo, Sebastian Gonzalez, Julián León, Pablo Tovar, Natalia Contreras and Ángela Rodriguez for their valuable comments on the early version of the manuscript and the mobile app.

